# LOX-1 and MMP-9 inhibition attenuates the detrimental effects of delayed rt-PA therapy and improves outcomes after acute ischemic stroke

**DOI:** 10.1101/2023.08.08.552551

**Authors:** Kajsa Arkelius, Trevor S Wendt, Henrik Andersson, Anaële Arnou, Michael Gottschalk, Rayna J Gonzales, Saema Ansar

## Abstract

**Background:** Acute ischemic stroke triggers endothelial activation that disrupts vascular integrity and increases hemorrhagic transformation leading to worsened stroke outcomes. Recombinant-tissue plasminogen activator (rt-PA) is an effective treatment; however, its use is limited due to a restricted time window and high risk for hemorrhagic transformation, which in part may involve activation of metalloproteinases (MMPs) mediated through lectin-like oxidized LDL receptor 1 (LOX-1). This study’s overall aim was to evaluate the therapeutic potential of novel MMP-9 and LOX-1 inhibitors in combination with rt-PA to improve stroke outcomes.

**Methods:** Thromboembolic rat stroke model was utilized to investigate the impact of rt-PA delivered 4h post-stroke onset as well as selective LOX-1 (BI-0115) and/or MMP-9 (JNJ0966) inhibitors given prior to rt-PA administration. Infarct size, perfusion, and hemorrhagic transformation were evaluated by MRI. Neurological function was assessed using sensorimotor functioning testing. Using an *in vitro*, human brain microvascular endothelial cell (HBMEC) model, cells were exposed to hypoxia plus glucose deprivation (3h)/reperfusion (12h) (HGD/R) and treated with rt-PA ± an MMP-9 and LOX-1 inhibition cocktail. MMP-9 activity was determined with zymography, and endothelial barrier marker gene expression and LOX-1 levels were evaluated via qRT-PCR and western blot respectively.

**Results:** Rt-PA treatment increased edema, hemorrhage, and worsened neurological outcomes post stroke. LOX-1 inhibition significantly improved neurological function and reduced edema after delayed rt-PA treatment. Hemorrhagic transformation, edema, and increased MMP-9 activity were attenuated by the MMP-9 inhibitor. Stroke induced increases in cerebrovascular LOX-1 expression correlated with increased MMP-9 activity and elevated activity correlated with increased edema, infarct volume, and decreased neurological function. In cultured HBMECs, LOX-1/MMP-9 inhibition differentially attenuated rt-PA-mediated increases in endothelial derived MMP-9 levels and activity, inflammation, and activation following HGD/R.

**Conclusion:** Here, we conclude that MMP-9/LOX-1 inhibition attenuates negative aspects of delayed rt-PA therapy leading to improved neurological function.

## Introduction

### Novelty and Significance

#### What is Known?

- Acute ischemic stroke (AIS) remains a global burden leading to significant mortality and morbidity rates to which there are only two currently FDA-approved therapies, including endovascular thrombectomy and administration of recombinant tissue plasminogen activator (rt-PA). However, these therapies exhibit an increased risk of hemorrhagic transformation leading to significant potential for worsened outcomes.
- Endothelium plays a critical role in maintaining the homeostatic environment of the blood-brain barrier (BBB) and when perturbed during AIS, can contribute significantly to the development and progression of stroke pathogenesis.
- Lectin-like oxidized low-density lipoprotein receptor (LOX-1) is a key contributor to the pathogenesis of vascular diseases such as atherosclerosis. Moreover, LOX-1 has been linked to matrix metalloproteinase 9 (MMP-9), an enzyme strongly associated with cerebrovascular pathogenesis during AIS.

#### What New Information Does This Article Contribute?

- We characterized the effects of acute ischemic injury, rt-PA administration, and LOX-1/MMP-9 inhibition on stroke outcomes using an in vivo as well as in vitro stroke model. We, in part, identified the impact of rt-PA on ischemic injury markers LOX-1, and MMP-9, and their role in the progression of AIS at the functional, neurological, and molecular levels.
- Using a thromboembolic stroke model, we characterized the spatiotemporal impact of ischemic stroke on MMP-9 activity within both the ipsilateral and contralateral hemispheric microvasculature and parenchymal tissues. Additionally, we identified a spatiotemporal dysregulation of the cerebrovascular and parenchymal tissue microenvironment which was differentially altered by rt-PA in the presence and absence of novel LOX-1 and MMP-9 inhibitors.
- We demonstrate links between LOX-1 and MMP-9 as contributors that dysregulate barrier integrity, increase inflammation, and lead to worsened neurological function. We also highlight the detrimental effects of rt-PA and the beneficial effects of selective LOX-1 and MMP-9 inhibitors at the level of the rodent cerebrovasculature and human brain microvascular endothelium. We propose that LOX-1 signaling leading to increased MMP-9 activity and neuroinflammation plays a critical role in worsened stroke pathogenesis.

Ischemic stroke affects over 9 million people each year worldwide, resulting in disability and death in 30% of the cases.^1^ AIS triggers a complex and multimodal response at the level of the cerebrovascular endothelium resulting in endothelial activation that, in part, disrupts vascular integrity setting off a cascade of secondary injuries that include barrier integrity loss, cerebrovascular inflammation, edema, hemorrhagic risk, infarct expansion, and worsened stroke outcomes.^2–5^ Effective therapies for AIS exist, such as thrombolysis and endovascular thrombectomy, which have improved the quality of life in stroke survivors.^6,7^ Recombinant tissue plasminogen activator (rt-PA) remains the only FDA-approved drug for AIS, however, due to its low efficacy, administration of rt-PA is restricted to a therapeutic window of 3-4.5h after stroke onset and has been shown to elicit an increased risk of hemorrhagic transformation and can lead to considerable morbidity.^6,8,9^

One of the underlying causes of increased hemorrhagic transformation risk appears to be due to the elevated levels of matrix metalloproteinase 9 (MMP-9),^10^ which has been shown to be directly activated by rt-PA.^11,12^ Intriguingly, acute hemorrhagic transformation <18-24h following stroke onset has been postulated to be leukocyte-derived MMP-9 and brain-derived matrix metalloproteinase 2 (MMP-2).^14^ However, the contributions of the cerebrovasculature to increased blood concentrations of MMP-9 and worsened outcomes during the acute phase of AIS clinically remains to be elucidated.

Several processes can induce an increase of MMP-9 expression, one being through the signaling pathway of the lectin-like oxidized low-density lipoprotein (oxLDL) receptor-1 (LOX-1), a receptor that is activated during hypoxia^15^ and has been well characterized in other cardiovascular pathologies.^16–19^ In previous studies we revealed a significant increase of LOX-1 6h after ischemic stroke in hypertensive male and female animals.^20,21^ LOX-1 was first characterized and cloned in vascular endothelial cells ^22^ and shown to induce endothelial cell dysfunction and activation. The connection between LOX-1 activation and MMP-9 production was established nearly a decade later in primary human aortic endothelial cells, which demonstrated that incubation with oxLDL induced an increase in levels of LOX-1, reactive oxygen species, NF-κB, and MMP-9 secretion.^23^

Although the LOX-1 receptor is capable of binding oxLDL, there are a multitude of other substrates that share very few structural similarities to oxLDL such as C reactive protein, fibronectin, and aged/apoptotic cells all of which are capable of binding to this scavenger receptor. ^24^ Therefore targeting LOX-1 may be beneficial in mitigating the pathogenesis of AIS, even in the absence of elevated oxLDL levels. Moreover, at the level of the cerebrovascular endothelium, LOX-1 signaling in part increases MMP-9 activity and decreases tight junction protein expression.^25^ A significant portion of the endothelium’s regulatory role in maintaining blood-brain barrier (BBB) homeostasis is attributed to specific proteins such as claudin-5,^26^ occludin,^27^ and intercellular adhesion molecule 1 (ICAM-1),^28,29^ which comprise critical aspects of endothelial cell tight junction function and extravasation. This further suggests that LOX-1 may be a potential therapeutic target to in part attenuate brain endothelial dyshomeostasis during AIS progression.

Studies have concomitantly investigated the effect of late rt-PA treatment and the involvement of LOX-1 and MMP-9 detrimental outcomes on BBB integrity.^30–32^ Thus, it is feasible to hypothesize that inhibition of MMP-9 and/or LOX-1 signaling during combination with rt-PA therapy could be a novel therapeutic approach to attenuate the increased risk of hemorrhagic transformation that accompanies rt-PA administration. Therefore, we hypothesized that combination LOX-1/MMP-9 inhibition would attenuate an ischemia/rt-PA-induced increase in MMP-9 activity, cerebrovascular permeability, and oxidative stress, which would result in improved neuroprotection and positive outcomes following ischemic stroke. In this study, we (i) determined the spatiotemporal dependent alteration in MMP-9 activation and LOX-1 levels within the cerebrovasculature and parenchymal tissues following stroke, (ii) characterized the potential detrimental impact of rt-PA on the cerebrovasculature and functional outcomes following stroke as well as assessed the beneficial effects LOX-1 and MMP-9 inhibition, both separately and when administered in combination, and (iii), directly assessed potential underlying mechanisms of cerebrovascular endothelial-derived MMP-9, integrity, and inflammation in response to ischemic injury, rt-PA, and combination therapy of LOX-1 and MMP-9 inhibition.

## Materials and Methods

The data supporting this study’s findings are available from the corresponding author upon reasonable request.

### Ethics

All procedures and animal experiments were performed in compliance with the European Community Council Directive (2010/63/EU) for Protection of Vertebrate Animals Used for Experimental and other Scientific Purposes guidelines. The ethical permit (Animal Inspectorate License No. 5.8.18-10593/2020) was approved by the Malmö-Lund Institutional Ethics Committee under the Swedish National Department of Agriculture.

### Thromboembolic stroke model and treatment

Performed as previously described, male Wistar rats (Janvier, France) were sedated with 3% isoflurane mixed in N_2_O/O_2_ (70:30) and then maintained at 1.5-2% for the duration of the surgery. ^33^ A heating pad connected to a rectal temperature probe assessed body temperature. The head was gently secured via a stereotactic device and a vertical cut midway between the right eye and ear allowed access for an incision in the temporal muscle. Next a craniectomy was performed and the middle cerebral artery (MCA) bifurcation was exposed after removing the dura. During the procedure, a laser doppler flow meter was secured on the skull within the right middle cerebral artery (MCA) region to monitor cerebral blood flow (CBFΑ-thrombin (12 UI Nordic Diagnostica AB, Sweden) was injected into the MCA lumen using a micropipette, which remained in place for 20min after the injection for clot stabilization. Surgery was deemed successful with a decrease in CBF of 70% that remained stable for one hour. Animals were randomized into the following groups: JNJ0966 ([N-(2-((2-methoxyphenyl)amino)-4’-methyl-[4,5’-bithiazol]-2’-yl)acetamide)(10mg/kg), BI-0115 (9-Chloro-5,11-dihydro-5-propyl-6H-pyrido[2,3-b][1,4]benzodiazepin-6-one)(10mg/kg), or the combination JNJ0966 plus BI-0115 i.p injection at 3.5h post occlusion at which rt-PA 3mg/kg (Alteplase, Boehringer Ingelheim AB, Germany) was given intravenously in the tail vein starting with a bolus dose of 10 % followed by a 40 min infusion The vehicle (saline) for rt-PA was administered at 4h after stroke induction. Animals were then re-anesthetized, and the laser Doppler placed in the same position as during the primary procedure to monitor CBF.

### Magnetic resonance imaging (MRI)

MRI was used to evaluate infarct lesion size, hemorrhage, and CBF at 24h post stroke onset. Animals were anesthetized with O_2_ mixed with 3 % isoflurane which then was lowered to 1.5-2 % for the duration of the imaging procedure. The animals’ breathing and body temperature were monitored during the procedure. Imaging was performed using a 9.4 T preclinical MRI horizontal bore scanner (Biospec 94/20, Bruker, Germany).

#### T2-weighted imaging

T2-weighted images were acquired using the RARE sequence: 25 axial slices, slice thickness 0.8mm, 256×146 matrix, in-plane resolution 137177 cl:140137µm^2^, TR = 270ms, TE = 33ms, bandwidth 33kHz, TA = 2min 25s.

#### T2*-mapping

T2*-maps were reconstructed from a multi gradient-echo sequence acquired with parameters as above except: TR = 1800ms, TE = 3.5ms to 58.5ms in steps of 5ms, bandwidth 69kHz, TA = 3 min 18s.

#### pCASL

Brain perfusion was measured using unbalanced Pseudo-continuous arterial spin labelling (pCASL) in PV6 using the implementation presented by Hirschler et al.^34^ The proposed protocol was used with small modifications as detailed below. Anatomical images were acquired with a 2D RARE sequence with TE 33ms, TR 2,5s, resolution 117185 cl:381117µm^2^, FOV 30185 cl:45230mm and slice thickness 0.8mm and with 23 slices and two averages. Two control and label and phase optimization prescans were acquired to determine the phase settings for label and control in the pCASL scans. Labelling was applied in the rat’s neck −2cm from the isocenter during 1.5s followed by a post-labelling delay of 300ms. Pulses and gradients for labelling were set as in the original publication. Readout was reported with a 2D EPI sequence with TE 13ms, TR 2 s, resolution 312185 cl:452313µm^2^, FOV 24185 cl:45230 mm and slice thickness 4 mm. Inversion efficiency was obtained as in the original publication with the exception that a resolution of 234 185 cl:452 234µm^2^ was used with 4 averages. The transmission coil was used in transmit-receive mode as reconstruction of arrayed coil data was not possible for the inversion efficiency measurement in the current implementation of the method. For the pCASL perfusion measurement the original implementation was followed with the modification of TE = 14.2ms, TR = 4s, resolution 234×234 µm^2^, FOV 18×29mm and slice thickness 2mm with 5 slices. Slices were placed to cover the stroke area otherwise centred around the isocentre. For the final T_1_ the modifications were TE=12.8ms and the geometry was taken from the preceding pCASL scan.

### Neurological evaluation

A 28-point neuroscore test and cylinder test prior to and 24h after stroke onset was used to assess neurological function. The 28-point neuroscore is a neurological composite test based on 11 different sensorimotor tasks. The animal’s performance of the various tasks was scored by an experienced evaluator blinded to the treatment groups. A score of 28-points, the accumulative score of the test, indicates a normal neurological function in a healthy rat. During the cylinder test rats are placed individually in a cylinder, and video recorded for 5min to evaluate the rats use of their left and right front paws when exploring the cylinder.^35^

### Tissue collection

Two different cohorts of animals were used, one for the temporal profile and the other for drug treatment. Animals used for the temporal profile were euthanized 3, 6, and 24h after stroke induction. Animals randomly selected for treatment were euthanized 24h post stroke after MRI and neurological function evaluation were performed. During tissue collection, rats were heavily sedated with 3 % isoflurane in a mixture of N_2_O/O_2_ (70:30) before intracardiac perfusion with saline was performed. Brains were collected and snap-frozen before being stored at −80°C until further use for western blot and zymography.

### Cerebral vessel and parenchyma isolations

The cerebral vasculature and parenchyma from the whole brain isolated from sham and stroke induced animals were separated according to a previously described method.^36^ Tissue from the infarct area in the cerebral cortex was dissected from the ipsilateral side, and the same area on the contralateral side was used as control. Dissected sections were homogenized in 0.1% 1M HEPES in Hank’s balanced salt solution (HBSS) and centrifuged at 2000g for 10min at 4°C.^36^ The resultant supernatant contained the parenchyma fraction. The pelleted vessels were resuspended in 18% dextran solution (18% dextran in 0.1% 1M HEPES/HBSS) and centrifuged at 4400g for 15min at 4°C. The supernatant was discarded, and the pellet was resuspended in 1% bovine serum albumin (BSA) in 0.1% 1M HEPES/HBSS and filtered twice with 1% BSA solution through a 20µm filter. The vessels on the filter were then suspended in 1% BSA solution and centrifuged at 2000g for 5min at 4°C.^36^ The supernatant was discarded, and the vessel fraction pellet was used immediately or frozen at −80°C until use.

### Human brain endothelial cell culture

Male primary human brain microvascular endothelial cells (HBMEC) were purchased from Cell Systems (Kirkland, WA, Catalogue number: ACBRI 376 Lot number: 376.05.02.01.2F) and cultured as previously described.^37^ Cells were received at *passage 3* and seeded in house at a density of 2 to 3 × 10^5^/cm^2^. HBMECs were grown in 5% CO_2_ and room air at 37°C in phenol red-free Complete Classic Medium (10% FBS; fetal bovine serum) supplemented with vendor specific guideline usage of Bac-off Antibiotic (Cell Systems, Catalogue number: 4Z0-643) and CultureBoost (Cell Systems, Catalogue number: 4CB-500). Monolayers were cultured in dishes coated with Attachment Factor (Cell Systems; Catalogue number: 4Z0-201), an extracellular matrix that served to enhance endothelial cell attachment, polarity, and cytoskeletal organization. Under these culture conditions, HBMECs reached 75-80% confluency within 4-5 days. Once cells reached confluency, selected plates were cryopreserved for future use, and the remaining HBMECs continued in culture and were studied at *passage 7*.

### Drug preparation and hypoxia plus glucose deprivation with reperfusion (HGD/R) model

All solutions were prepared fresh under sterile conditions on the same day of administration for planned experiments. Human tissue plasminogen activator (rt-PA) (Fisher Scientific; Catalogue number: 612200100UG), was prepared to a stock solution (1mg/mL) in ddH_2_O and administered at a concentration of 12.5µg/mL. Similarly, JNJ0966 (MedChemExpress; Catalogue number: HY-103482), was prepared to a stock solution (10mM) in dimethyl sulfoxide (DMSO) and administered at a concentration of 5µM. BI-0115 (Boehringer Ingelheim; Catalogue number: BI-0115), was prepared to a stock solution (10mM) in dimethyl sulfoxide (DMSO) administered at a concentration of 10µM. Vehicle solutions utilized in the control groups were generated in tandem and consisted of a DMSO/DPBS solution prepared at a non-toxic concentration (DMSO <0.1%). These solutions were maintained at 37°C during preparation and administration to HBMEC cultures. Normoxic control plates containing Complete Classic Medium were replaced with fresh medium and incubated in a separate CO_2_ incubator at 37°C at 21% O_2_. Plates designated for HGD/R exposure, had the medium replaced with modified DMEM (ThermoFisher Scientific; Catalogue number: A14430-01) media containing no L-glutamine, phenol red, sodium pyruvate, or D-glucose and then placed in a hypoxic sub chamber (BioSpherix) at 37°C and flushed with a medical grade gas mixture of 1% O_2_, 5% CO_2_, and nitrogen balance and exposed for 3h. Next, some plates were returned to 21% O_2_, received fresh Complete Classic Medium, and were immediately treated with vehicle, rt-PA, or rt-PA + JNJ + BI. Following treatment with select compounds they were then incubated in a CO_2_ incubator at 37°C at 21% O_2_ for 24h. Designated plates for temporal studies were either collected immediately following 3h of HGD exposure or received fresh Complete Classic Medium and were then incubated in a CO_2_ incubator at 37°C at 21% O_2_ for 12 and 24h.

### Quantitative real time PCR

Quantitative real time PCR (qRT-PCR) was utilized to measure changes in mRNA levels of *MMP-9*, *TIMP1*, *IL-1β*, *CLDN5*, *OCLN*, *ICAM1*, *SOD1*, and *TJP1*. Briefly, RNA was extracted using the Qiagen kit (ThermoFisher, Cat. No. 74034) according to the manufacturer’s instructions. Purity and concentration of extracted RNA was confirmed using a Nanodrop 2000 (ThermoFisher Scientific). RNA was reverse transcribed to make cDNA using the SuperScript III First-Strand Synthesis System (ThermoFisher Scientific; Catalogue number: 18080051) according to the manufacturer’s instructions. All primer efficiencies were assessed via serial dilution from 200ng/L to 0.02ng/L and were found to be within 90-110% efficient. Within each well a mixture of cDNA (5µL), target primers diluted in DEPC (5µL; 0.4µM forward primer, 0.4µM reverse primer), and POWER SYBR Green PCR Master Mix (ThermoFisher Scientific; Catalogue number: 4368706) was added in order. The 96-well plates were then centrifuged (Thermo Sorvall Legend RT+) at 1,000g for 2 min and then immediately loaded into the QuantStudio 6 and 7 Flex real-time PCR system (ThermoFisher Scientific). A two-step cycling protocol was implemented to collect cycle threshold (Ct) values of specific target amplicons and housekeeping gene *GAPDH* using the ΔΔCt relative quantification setting within QuantStudio 6. The reaction involved 2min at 50°C, an initial denaturing step at 95°C for 10min followed by 40 cycles of 95°C (denature) for 15s and primer-specific annealing temperature for 1min. Primer annealing temperatures for *MMP-9*, *TIMP1*, *IL-1β*, *CLDN5*, *OCLN*, *ICAM1*, *SOD1*, *TJP1*, and *GAPDH* are 64.1C, 67.4°C, 64.4°C, 68.0°C, 65.7°C, 68.2°C, 62.4°C, 65.6°C, and 64.4°C respectively. The primer sequences used are described in **Supplemental Table 1**.

### Protein extraction and concentration determination

#### Parenchyma and vessels (in vivo study)

Parenchyma and cerebral vascular fractions were homogenized in lysis buffer containing protease and phosphatase inhibitors (ThermoScientific #FNN 0091, SIGMA # p2850, #p8340) and incubated on ice for 20min. Parenchyma fractions were centrifuged at 20000g for 20min at 4°C, and cerebral vasculature fractions at 10000g for 10min at 4°C, subsequently supernatant was collected. Total protein concentration was determined using a Bio-Rad protein assay kit measuring absorbance at 595nm on a Bio-Rad microplate reader. Lysates were stored at −80°C until further use for western blot and zymography.

#### Conditioned Media (in vitro study)

Media was extracted from cultures following exposure/treatment and centrifuged at 1,000g for 5min to remove any cellular debris and the supernatant was removed and utilized as conditioned media. Total protein within conditioned media was determined on the day of zymography experimentation and determined by using a bicinchoninic acid protein assay kit (ThermoFisher Scientific) according to the manufacturer’s protocol and measured on a Safire II (Tecan) plate reader.

### Western blot

Proteins of interest were evaluated in vessels and parenchyma fractions. Samples were prepared with Laemmli sample buffer and boiled at 95°C for 4min. Molecular weight marker PageRuler Plus Prestained Protein Ladder ThermoFisher) was used to determine the protein of interest. Proteins were then transferred to a nitrocellulose membrane (Bio-Rad) blocked with 5% milk/Tween-20-TBS (T-TBS) 0,1%. Membranes were incubated with primary antibodies, LOX-1 (1:1000 #ab60178 Abcam), Occludin (1:500 #71-500 Invitrogen), VCAM-1 (1:500 #sc-13160 Santa Cruz Biotechnology), diluted in 5% bovine serum albumin (BSA) T-TBS 0,1%, overnight at 4°C. Membranes were washed 3 × 5min in T-TBS 0,1%. Subsequently, the membranes were incubated with the corresponding secondary antibody, anti-mouse (1:2000 – #7076 Cell Signaling), or anti-rabbit (1:2000 – #7074 Cell Signaling) for 1h at RT. After that, membranes were washed 4 × 5min in T-TBS 0,1% and 5min in TBS. Membranes were developed with SuperSignal™ West Dura Extended Duration Substrate (#34075 ThermoFisher) for 5min in a dark room. Bands are visualized by FujiLAS1000 Luminescent Image Analyzer (Stamford, CT, USA). Levels of ß-actin (1:50 000, Sigma Aldrich, #A3854) were used as the loading control.

### Zymography

Conditioned media, parenchyma, and vessel samples (20µg) were diluted in Tris-glycine SDS sample buffer and the equally diluted samples and fluorescent standards (LI-COR Biosciences) were loaded onto Novex 10% zymogram plus gelatin protein gels (ThermoFisher Scientific; Catalogue number: ZY00100BOX). Protein separation occurred by SDS-PAGE using a mini-PROTEAN Tetra electrophoresis system (Bio-Rad Laboratories) at 100V for 2h and held at 4°C. After electrophoresis, the gel was incubated in zymogram renaturing buffer (ThermoFisher Scientific; Catalogue number: LC2670) for 0.5h at RT with gentle agitation. Zymogram renaturing buffer was decanted and replaced with zymogram developing buffer (ThermoFisher Scientific; Catalogue number: LC2671) and allowed to equilibrate the gel for 0.5h at RT with gentle agitation. Following equilibration, developing buffer was decanted and gels washed with nano-pure H_2_O and replaced with fresh 1x zymogram developing buffer and developed for either 24h (conditioned media) or five days (parenchyma and vessel isolates) at 37°C. After developing, gels were briefly washed with nano-pure H_2_O and stained with Coomassie blue staining solution (2.5g Coomassie Brilliant blue R-250, 450ml 100% methanol, 100ml 100% acetic acid, 400ml nano-pure H_2_O) for 2h at RT with gentle agitation. Gels were then washed with nano-pure H_2_O and de-stained with Coomassie blue de-staining solution (450ml 100% methanol, 100ml 100% acetic acid, 400ml nano-pure H_2_O) for 20min at RT with gentle agitation. The area of gelatin degradation at approximately 82kDa and 63kDa was indicative of MMP-9 and MMP-2 activity respectively and appeared as distinct white bands. The intensities of both MMP-9 and MMP-2 in conditioned media were obtained by an Odyssey Classic infrared imager (LI-COR) and inverse band densities were then analyzed using ImageStudio 3.0 software (LI-COR). For parenchyma and vessel isolates, gels were imaged using the ChemiDoc XRS+ (BIO-RAD), and band intensity analyzed in ImageJ.

### Extracellular H_2_O_2_ Quantification

Conditioned media (CM) was obtained as previously described in *condition media* section and utilized to ascertain concentration of hydrogen peroxide (H_2_O_2_). The aqueous pierce quantitative peroxide assay kit (ThermoFisher Scientific; Catalogue number: 23280) was utilized per manufacturer’s instructions to measure levels of H_2_O_2_ across treatment groups. Absorbance was then measured on a Safire II (Tecan) plate reader at 595nm and colorimetric quantification of H_2_O_2_ was compared to standards. Each sample was run in duplicate.

### Statistical Analyses

The number of animals and independent HBMEC samples are referred to as “*n*” and P < 0.05 was considered statistically significant. Statistical analyses were performed using GraphPad Prism version 9.3.0 for Windows, GraphPad Software, USA, www.graphpad.com. In vitro: Data is presented with mean±SD. Comparisons between multiple groups were made using one-way ANOVA with Tukey’s multiple comparisons post-hoc test, and direct comparisons with unpaired t-test. In vivo: Data are presented with mean±SD. Comparisons were made in either the ipsilateral hemisphere or the contralateral hemisphere using one-way ANOVA with Tukey’s multiple comparisons post-hoc test.

## Results

### Spatiotemporal alterations in MMP-9 activity correlate with LOX-1 expression following thromboembolic stroke

Cerebral vascular and parenchymal spatiotemporal MMP-9 activity was measured via a zymography assay at 3, 6, and 24h (**Figs. 1A**) following the onset of experimental thromboembolic stroke. MMP9 activity is illustrated in brain vessels (**Figs. 1B-C)** and parenchymal tissues (**Figs. 1D-E)** from contralateral and ipsilateral hemispheres. We observed a spatiotemporal increase in both vascular and parenchymal MMP-9 activity at 6h post stroke onset that was specific to the ipsilateral hemisphere. Since LOX-1 has been well investigated in the context of vascular diseases such as atherosclerosis,^40^ hypertension,^19^ and stroke^20^, we hypothesized that LOX-1 levels may correlate with increased MMP-9 activity with the potential to lead to worsened outcomes via increased endothelial dysfunction and BBB permeability (**Fig. 1F**). Although we did not observe changes in LOX-1 expression levels within the cerebral vasculature and parenchymal tissues following thromboembolic stroke (**Supplemental Fig. 1**), we did note that within individual rats, there was a significant correlation between increased LOX-1 expression and MMP-9 activity within the cerebral vasculature in the ipsilateral hemisphere of injury (**Fig. 1G**).

**Figure 1.**
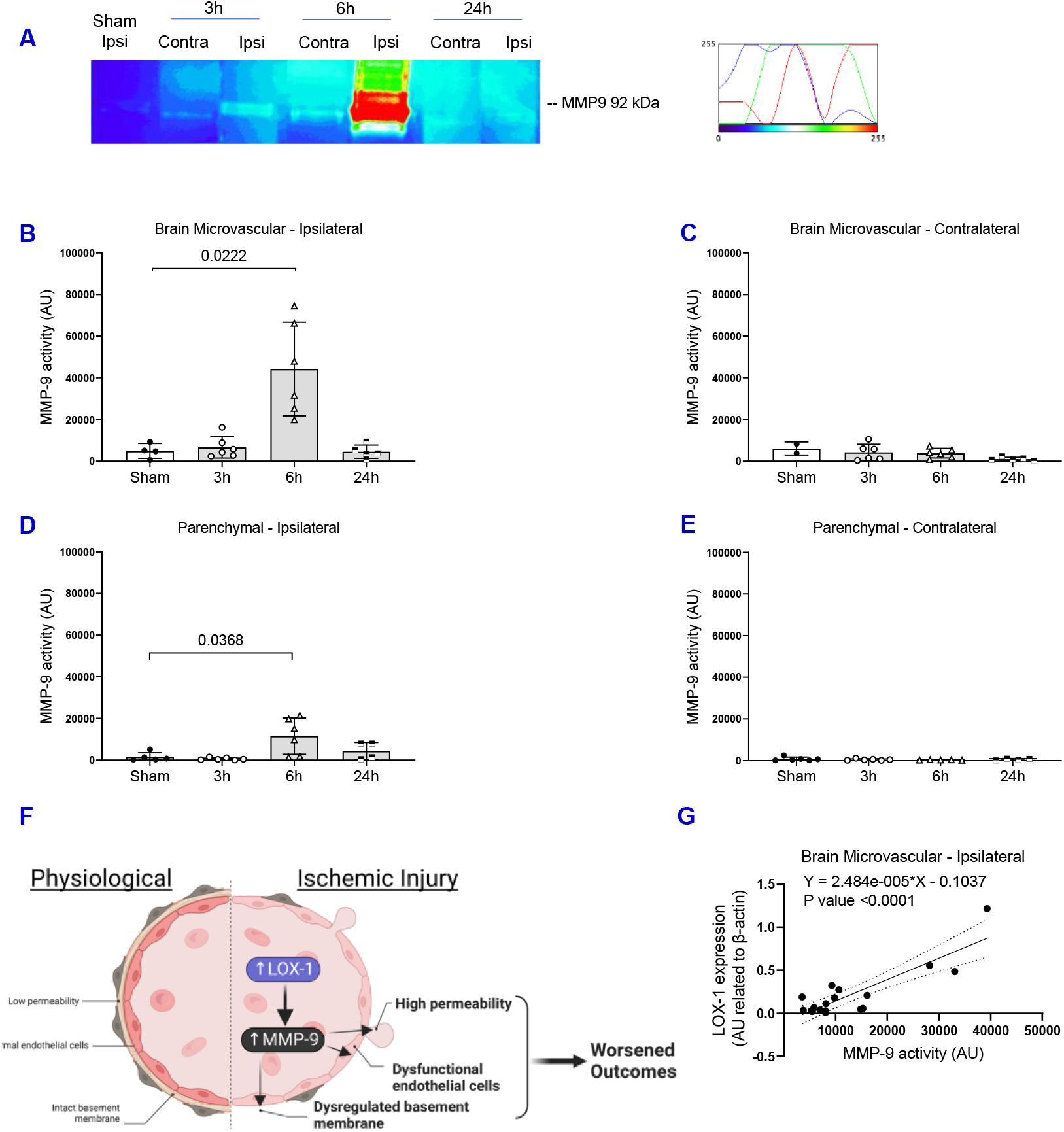
Spatiotemporal brain microvascular and parenchymal MMP-9 protein activity following thromboembolic stroke with as correlation to brain microvascular LOX-1 protein expression. (A) Representative gelatin zymography illustrating the band migration and enzymatic activity for matrix metalloproteinase 9, MMP-9 (92kDa), in isolated brain microvascular tissue from the ipsilateral and contralateral hemispheres following either sham or thromboembolic stroke at 3, 6, and 24h. Bar graphs depict MMP-9 activity within isolated microvessels (B-C) and parenchymal tissue (D-E) from the ipsilateral (B-D) and contralateral (C-E) hemispheres relative to injury by inverse densiometric analysis and expressed as arbitrary units (AU) *±* SD. *n* = 2-6 animals (data are mean ± SD). (F) Graphical illustration of our hypothesis demonstrating the linkage between increased levels of LOX-1 and MMP-9, which lead to worsened outcomes by mediating dysfunctional endothelium, increased permeability, and dysregulating the basement membrane. (G) Graph depicting the levels of LOX-1 protein normalized to β-actin (y-axis) and the corresponding MMP-9 activity (x-axis) observed in brain microvessels isolated from the ipsilateral hemisphere of rats at 24h following either sham or stroke. *n* = 18 animals. Linear regression analysis

### Selective MMP-9 and LOX-1 inhibition in combination with delayed rt-PA therapy attenuated vascular MMP-9 activity, reduced hemorrhagic transformation, decreased infarct size, and improved neurological function

We next examined the impact of selective MMP-9 and LOX-1 inhibition using JNJ0966 and BI0115 respectively, on functional outcomes following thromboembolic stroke. The safety of delayed rt-PA in the presence of a single drug treatment or in combination were examined by evaluating infarct volume, edema percentage, hemorrhagic incidence, diffusion, and perfusion using MRI. Representative MRIs are illustrated in **Fig 2A**. Similar to the clinical scenario, rt-PA in the thromboembolic model increased the risk of edema (**Fig. 2C**) and hemorrhagic transformation (**Fig. 2D**) percentage at 24h post-stroke induction, which was attenuated by LOX-1 and MMP-9 inhibition. We did not observe an increase in infarct volume with rt-PA compared to saline, however when combining rt-PA treatment with the LOX-1 and/or the MMP-9 inhibition there was a trend towards limiting the formation of large infarct lesions compared to saline or rt-PA treated animals (**Fig. 2B**). Moreover, while hemorrhages were present in animals treated with the MMP-9 and LOX-1 inhibitors alone or in combination, the number and size of the hemorrhages were relatively smaller in comparison to the rt-PA treated animals (**Fig. 2D**). Perfusion within the ipsilateral hemisphere was restored in all saline-treated animals 24h after stroke onset, except for animals treated with the rt-PA and MMP-9 inhibitor (**Fig. 2E**). In terms of neurological function, we observed that following thromboembolic stroke, animals treated with saline, rt-PA, and rt-PA + MMP-9 inhibitor had reduced functionality in their contralateral forepaw at 24h (**Fig. 2F**). These data together suggest that MMP-9 and/or LOX-1 inhibition may provide protective effects at the level of the BBB following an acute ischemic injury. However, the animals treated with the LOX-1 inhibition either alone or in combination with the MMP-9 inhibitor conjugated with rt-PA exhibited forepaw utilization similar to the sham-operated animals (**Fig. 2F**). Comparable results were also observed in the broader sensorimotor test of neurological function (28-point neurological score). While delayed rt-PA therapy appeared to have further detrimental effects on stroke outcome compared to saline-treated animals, LOX-1 inhibition significantly improved neurological function in animals treated with rt-PA (**Fig. 2G**). Moreover, combination of both the LOX-1 and MMP-9 inhibitors showed a similar trend of improvement compared to treatment with the LOX-1 inhibitor alone. Together these data suggest that targeting LOX-1 and MMP-9 following a thromboembolic stroke is in part acting at the level of the BBB is neuroprotective. We next assessed vascular derived MMP-9 activity and LOX-1 expression within the ipsilateral hemisphere. We observed in animals treated with saline at 24h post-stroke there was no increase in MMP-9 activity within the vasculature both within the ipsilateral (**Fig. 2H**) and contralateral (**Supplemental Fig. 2A**) hemispheres relative to sham. However, the administration of rt-PA following the induction of stroke increased vascular derived MMP-9 activity within the ipsilateral hemisphere compared to sham and was attenuated by the MMP-9 inhibitor (**Fig. 2H**). Concomitantly, rt-PA decreased vascular derived MMP-9 activity in the contralateral hemisphere compared to sham and activity was not altered by the MMP-9 inhibitor, JNJ0966 and/or the LOX-1 inhibitor, BI-0115 (**Supplemental Fig. 2A**). These data together suggest that at 24h post-stroke the delayed administration of rt-PA spatially increased MMP-9 activity within the ipsilateral hemisphere of injury which clinically correlates to an increased risk of hemorrhagic transformation observed previously. Moreover, the selective MMP-9 inhibitor, JNJ0966, significantly reduced the rt-PA mediated increase in MMP-9 activity which could suggest it plays a protective effect on BBB integrity following stroke. In terms of addressing LOX-1 expression, we observed that relative to saline injection, the administration of rt-PA alone resulted in an increase in LOX-1 levels in the vasculature within the ipsilateral hemisphere (**Fig. 2I**). Moreover, treatment with the MMP-9 and/or LOX-1 inhibitor(s) attenuated the rt-PA mediated increase in LOX-1 protein expression (**Fig. 2I**). These data further support the positive correlation analysis of LOX-1 protein and MMP-9 activity levels within the vasculature, suggesting a detrimental link between LOX-1 and MMP-9 following AIS.

**Figure 2.**
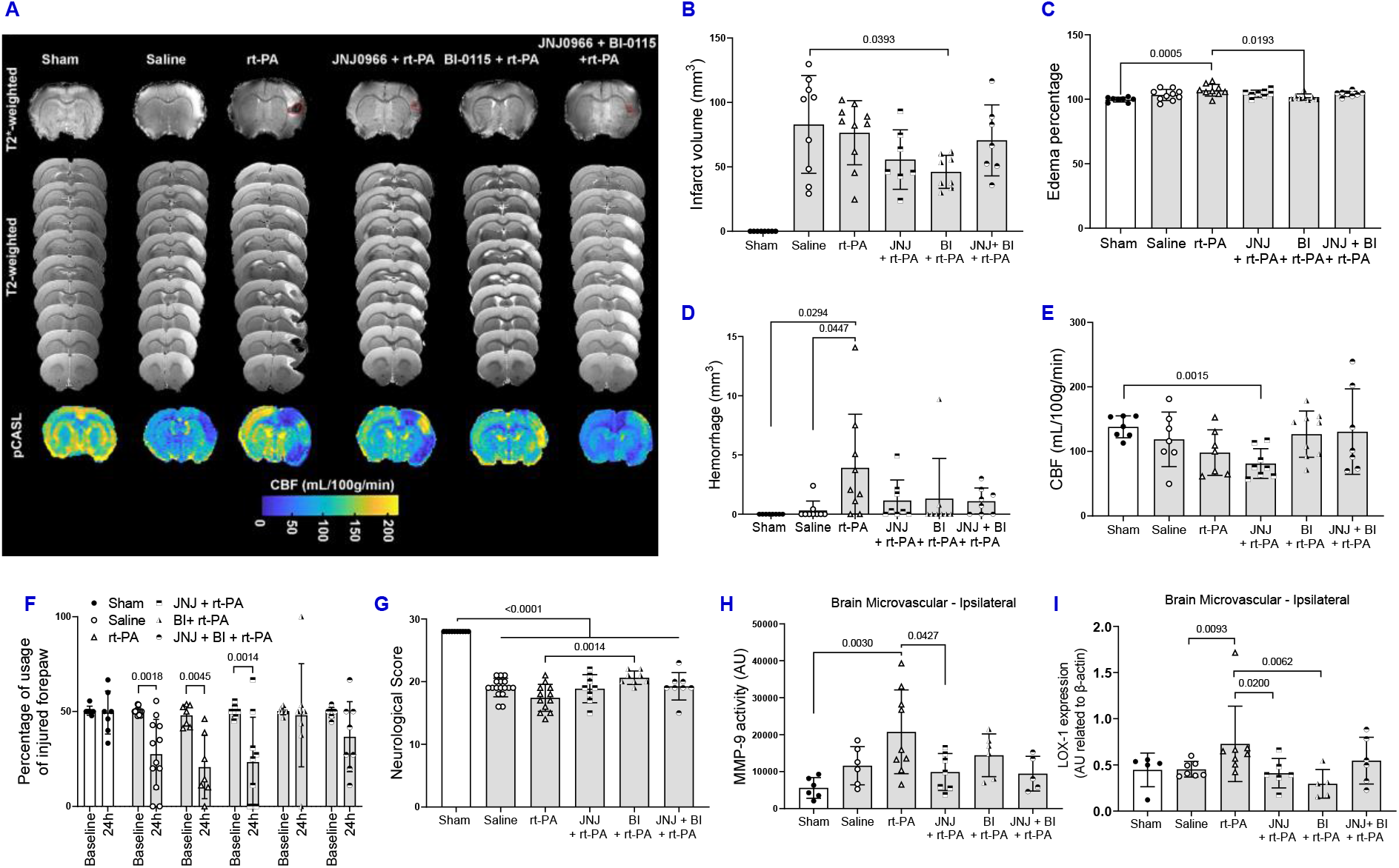
Impact of rt-PA +/− LOX-1 and/or MMP-9 inhibition following thromboembolic stroke on infarct, edema, hemorrhage, cerebral blood flow, neurological function, MMP-9 activity, and LOX-1 expression. (A) Representative T2- and T2*-weighted MRI as well as pCASL images of rat brains at 24h reperfusion and ischemic post injury. (B) Bar graph depicts infarct volume as measured by MRI analysis and volume expressed as millimeters^3^ (mm^3^) *±* SD. *n* = 7-10 animals. (C) Bar graph depicts edema as measured by MRI analysis and expressed as the percentage change from sham *±* SD. *n* = 8-10 animals. (D) Bar graph depicts hemorrhage as measured by MRI analysis and volume expressed as millimeters^3^ (mm^3^) *±* SD. *n* = 8-9 animals. (E) Bar graph depicts cerebral blood flow (CBF) as measured by MRI analysis and expressed mL/100g/min *±* SD. *n* = 6-8 animals. (F) Bar graph depicting the percentage of injured paw usage measured prior to (Baseline) and at 24h following treatment. *n* = 7-12 animals. (G) Bar graph depicts neurological score measured by blinded analysis and expressed as a scale between 0-28 with 0 representing the most severe and 28 representing the least severe. *n* = 8-17 animals. (H) Bar depicts MMP-9 activity within isolated microvessels from the ipsilateral hemisphere relative to injury by inverse densiometric analysis and expressed as arbitrary units (AU) *±* SD. *n* = 6-9 animals (I) Bar graph illustrates densiometric analysis of LOX-1 protein levels expressed as arbitrary units (AU) normalized to β-actin *±* SD. *n* = 5-9 animals.

### Temporal effect of ischemic-like injury on MMP-9 activity and barrier integrity in human brain microvascular endothelial cells

Utilizing primary human brain endothelial cells (HBMECs) we assessed mRNA levels of various markers of inflammation and barrier integrity following 3h of HGD and during 12 and 24h reperfusion (HGD/R) (**Fig. 3A**). We concomitantly observed an increase in MMP-9 and decrease in TIMP1 mRNA levels in HBMECs following 3h HGD exposure as well as at 24h HGD/R (**Figs. 3B & C**). We observed an increase in IL-1β mRNA levels at 3h HGD and during reperfusion levels decreased back to expression levels observed during normoxia (**Fig. 3D**). Counter to our hypothesis there was an initial increase in CLDN5 mRNA following 3h of HGD and a decrease in mRNA levels during simulated reperfusion (**Fig. 3E**). In comparison to CLDN5 levels, there was a concomitant increase in occludin mRNA at 24h post simulated reperfusion (**Fig. 3F**) which may suggest at a transcriptional level there is a preferential switch to occludin rather than claudin-5 expression following simulated ischemic injury. Similarly, we observed an initial decrease in ICAM1 mRNA at 3hHGDexposure with no relative change at 12 or 24h post HGD and during reperfusion (**Fig. 3G**). These data suggest that at the level of the human cerebrovascular endothelium there is a temporal effect of ischemia-reperfusion on mechanisms to increase MMP-9 activity and decrease endothelial barrier integrity.

**Figure 3.**
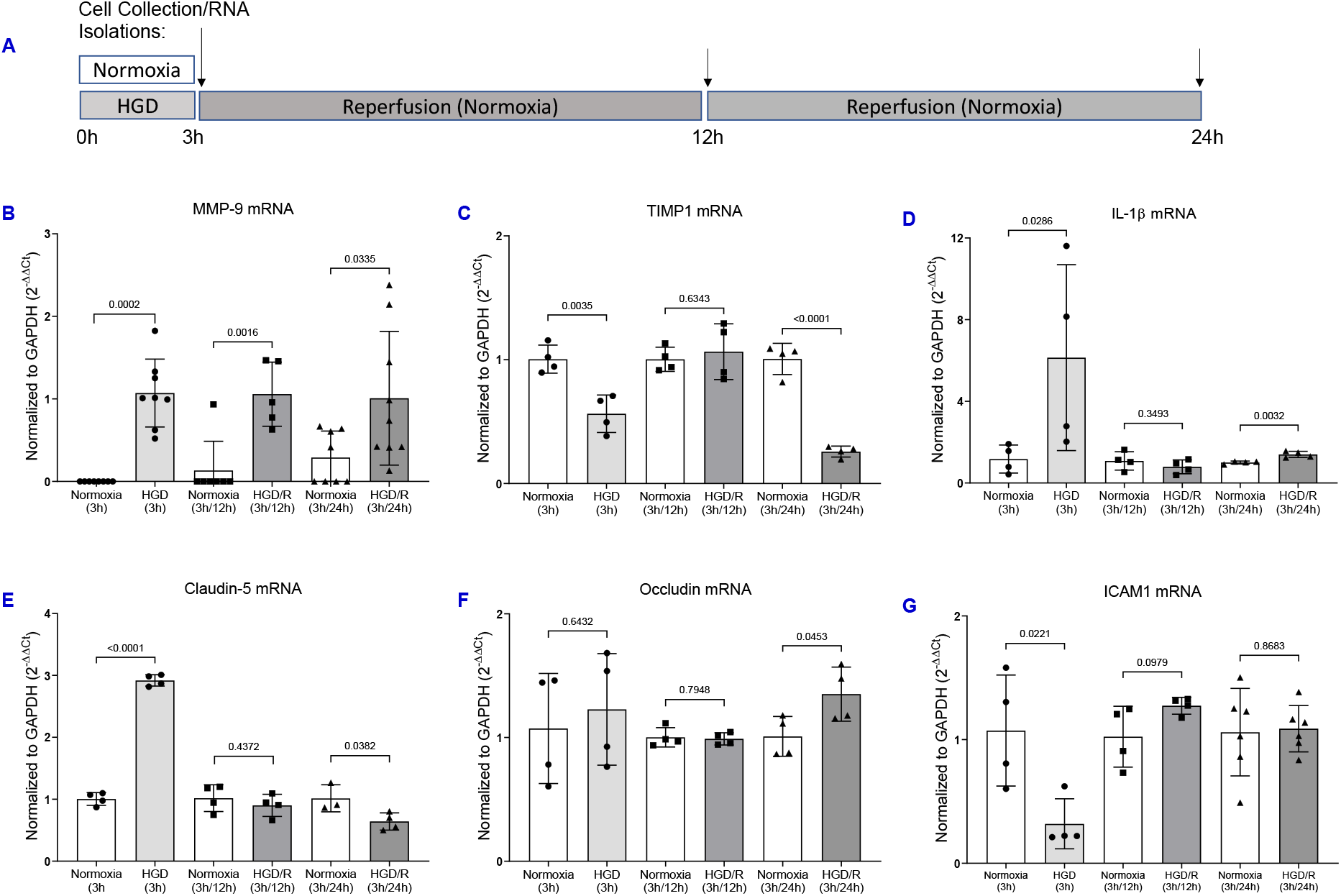
Temporal effect of in vitro ischemia/reperfusion injury on primary human brain microvascular endothelial cell (HBMEC) MMP-9, inflammatory, and barrier markers. (A) Illustration depicts the treatment timeline of in vitro ischemia/reperfusion injury on HBMECs exposed to either normoxia or HGD for 3h followed by either immediate collection or collection at 12 and 24h post reperfusion. (B-G) Bar graph depicts qRT-PCR mediated quantitation of MMP-9 (B), TIMP-1 (C), IL-1b (D), claudin-5 (E), Occludin (F), and ICAM-1 (G) mRNA expression normalized to the housekeeping gene (GAPDH) and expressed as 2^−ΔΔ*ct*^. Samples were run in duplicate (data are mean ± SD). *n* = 4-9 independent samples.

### Endothelial derived MMP-9 and MMP-2 activity is differentially increased by rt-PA following HGD/R injury

Zymography of conditioned media revealed two distinct bands at approximately 93kDa and 65kDa, corresponding to MMP-9 and MMP-2 respectively (**Figs. 4A & D**). We first assessed the impact of HGD/R on HBMEC derived MMP-9 activity, which in concordance with our observations in saline treated mice (**Fig. 2H**), was not different following in vitro HGD/R injury (**Fig. 4A**). Similarly, extracellular HBMEC derived MMP-2 activity was not different following HGD/R. (**Fig. 4B**). We next examined the effects of rt-PA +/− MMP-9/LOX-1 inhibition on MMP-9 activity, we observed that rt-PA therapy did not alter HBMEC derived MMP-9 activity, although we did detect a trend increase in some of the HBMECs treated with rt-PA (**Fig. 4C**). While there was no difference in HBMEC derived MMP-9 activity with rt-PA treatment, there was a significant increase in MMP-2 activity at HGD 3h /R 24h / with rt-PA compared to vehicle (**Fig. 4D**).

**Figure 4.**
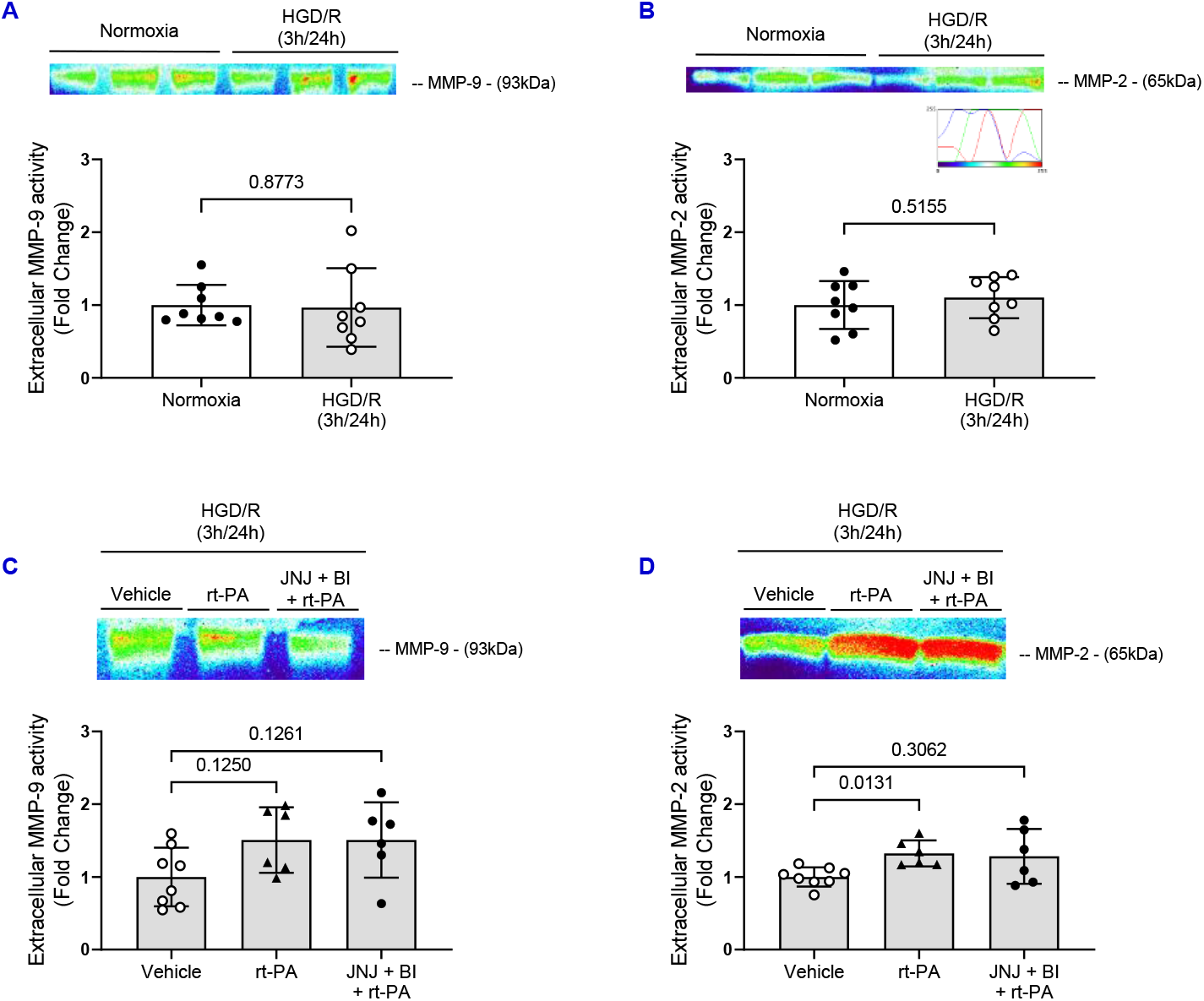
Impact of HGD/R +/− rt-PA +/− LOX-1 and MMP-9 inhibition on HBMEC MMP-9 and MMP-2 activity. (A-D) Representative gelatin zymography gels illustrating the band migration and enzymatic activity for two distinct matrix metalloproteinase bands, MMP-9 (93kDa) and MMP-2 (65kDa), in conditioned media collected from HBMECs. Bar graphs depict extracellular MMP-9 (B & E) and MMP-2 (C & F) activity by inverse densiometric analysis and expressed as fold change ± SD. *n* = 8 independent samples.

### MMP-9/LOX-1 inhibition in combination with rt-PA elicits differential effects on HBMEC MMP-9, inflammation, tight junction, and adhesion molecule mRNA expression

To next address potential mechanisms promoting cerebrovascular permeability following an ischemic injury and in the context of rt-PA, we assessed mRNA levels of multiple mediators in our HBMEC model following 3h of HGD and 24h reperfusion in the presence or absence of MMP-9/LOX-1 inhibition (**Fig. 5A**). HGD 3h/R 24h increased MMP-9 mRNA expression which was attenuated with rt-PA therapy alone or combination with MMP-9/LOX-1 inhibitors (**Fig. 5B**). Comparable to the observed MMP-9 response, administration of rt-PA alone did not alter TIMP1 mRNA levels, but when treated MMP-9/LOX-1 inhibition and rt-PA there was a significant increase in TIMP1 mRNA levels (**Fig. 5C**). We next assessed levels of HBMEC IL-1β mRNA and found that treatment with rt-PA significantly increased IL-1β mRNA at HGD 3h/R 24h (**Fig. 5D**). Concomitantly, the co-therapy of MMP-9/LOX-1 inhibition with rt-PA attenuated the effects of rt-PA alone and reduced levels of IL-1β mRNA (**Fig. 5D**). Additionally, we measured levels of extracellular hydrogen peroxide (H_2_O_2_) to further assess potential mechanisms behind MMP-9 activation and observed that HGD/R increased levels of HBMEC derived extracellular H_2_O_2_ when compared to normoxic controls, but rt-PA alone or in combination with MMP-9/LOX-1 inhibition did not alter these levels (**Supplemental Figs. 3A & B**). Moreover, we observed no difference in the levels of SOD1 mRNA under any exposure/treatment conditions (**Supplemental Figs. 3C & D**). Overall, it appears that combination therapy of MMP-9/LOX-1 inhibition with rt-PA could promote protective effects on HBMEC derived MMP-9 and inflammation at HGD 3h/R 24h.

**Figure 5.**
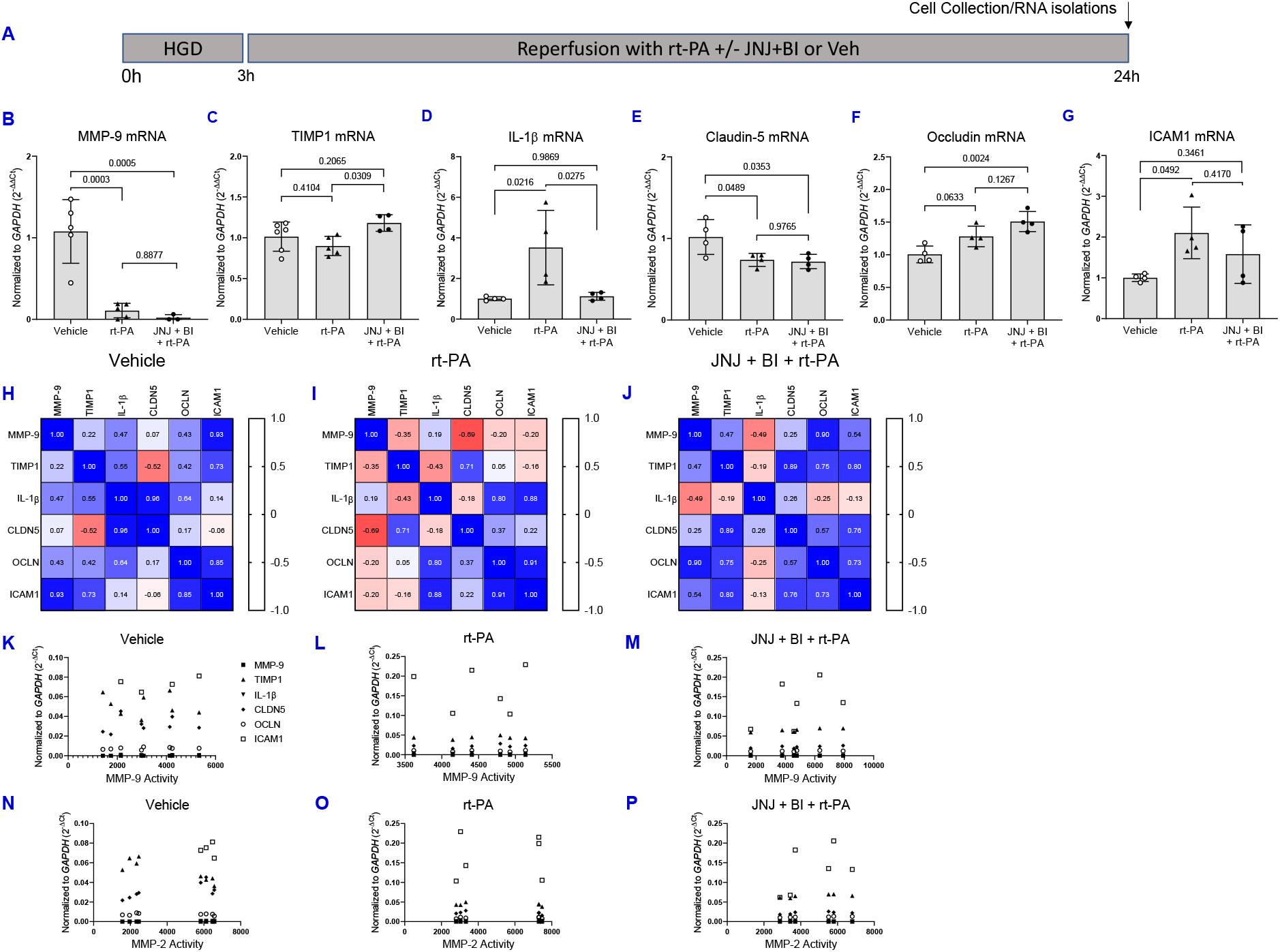
Impact of rt-PA +/− LOX-1 and MMP-9 inhibition on HBMEC MMP-9, inflammatory, and barrier markers following in vitro ischemia/reperfusion injury. (A) Illustration depicts the treatment timeline. (B-G) Bar graphs depict qRT-PCR mediated quantitation of MMP-9 (B), TIMP-1 (C), IL-1β (D), Claudin-5 (E), Occludin (F), and ICAM-1 (G) mRNA normalized to the housekeeping gene (GAPDH) and expressed as 2^−ΔΔ*ct*^. *n* = 3-6 independent samples. (H-J) Graphical representation of Pearson correlation matrices of HBMEC mRNA levels on a scale from −1 to 1 following HGD/R and either treated with vehicle (H), rt-PA (I), or JNJ + BI + rt-PA (I). *n* = 4-6 independent samples. (K-P) Graphical representation of Pearson correlation of extracellular HBMEC MMP-9 or MMP-2 activity against mRNA levels following HGD/R and either treated with vehicle (K & N), rt-PA (L & O), or JNJ + BI + rt-PA (M & P). (K-P) *n* = 4-8 independent samples.

To assess the underpinnings of endothelial integrity in the context of rt-PA and whether MMP-9/LOX-1 inhibition alters this response, we measured mRNA expression of tight junctions and adhesion molecule markers during HGD 3h/R 24h. We observed decreased CLDN5 mRNA levels following rt-PA therapy alone or in the presence of MMP-9/LOX-1 inhibition (**Fig. 5E**). In contrast to CLDN5 mRNA expression, there was an increase in OCLN mRNA in HBMECs treated with the combination of MMP-9/LOX-1 inhibition plus rt-PA therapy compared to vehicle (**Fig. 5F**). Thus suggesting, at 3h/24h HGD/R there might be a preferential switch within HBMECs towards occludin expression instead of claudin-5. This is further supported by our observations that HGD/R decreased mRNA levels of zonula occludens 1 (TJP1) (**Supplemental Fig. 3E**), which is critical for the stabilization of claudin-5 at the BBB. [Reviewed in ^41^] Moreover, we observed an increase in HBMEC ICAM1 mRNA expression following rt-PA therapy compared to HGD/R alone (**Fig. 5G**). In efforts to identify concomitant effects of HGD/R on mRNA in the presence or absence of rt-PA alone or in combination with MMP-9/LOX-1 inhibition we performed Pearson correlation analysis (**Figs. 5H-J**). We next sought to assess the correlations between each mRNA targets with the observed extracellular HBMEC MMP-9 and MMP-2 enzyme activity following HGD/R in the presence or absence of rt-PA alone or in combination with MMP-9/LOX-1 inhibition (**Figs. 5K-P**). Breakdowns of each correlation can be found in **Supplemental Table 1**. Together, these data suggest that HGD/R mediates a potential decrease in endothelial barrier integrity via a decrease CLDN5 and TJP1 mRNA. Furthermore, rt-PA therapy may exacerbate this barrier dysfunction by differentially altering CLDN5 and OCLN and increasing ICAM1 mRNA levels, which is partially attenuated by the combination therapy of MMP-9/LOX-1 inhibition with rt-PA.

### MMP-9 activity is correlated with worsened stroke outcomes

In efforts to further elucidate the detrimental role of MMP-9 in the pathogenesis of stroke outcomes such as edema, infarct volume, and neurological function we performed correlative analyses of vascular MMP-9 activity with these outcomes across all treatment groups. The individual treatment groups can be found in **Table 1**. Following correlative analysis, we determined that increased MMP-9 activity at 24h-post thromboembolic stroke was significantly correlated with increased edema (**Fig. 6A**), worsened neurological function (**Fig. 6B**), and increased infarct volume (**Fig. 6C**). These data suggest that targeting of vascular derived MMP-9 either by direct steric inhibition or potentially through the combination therapy of LOX-1 and MMP-9 inhibition could lead to improved stroke outcomes.

**Figure 6.**
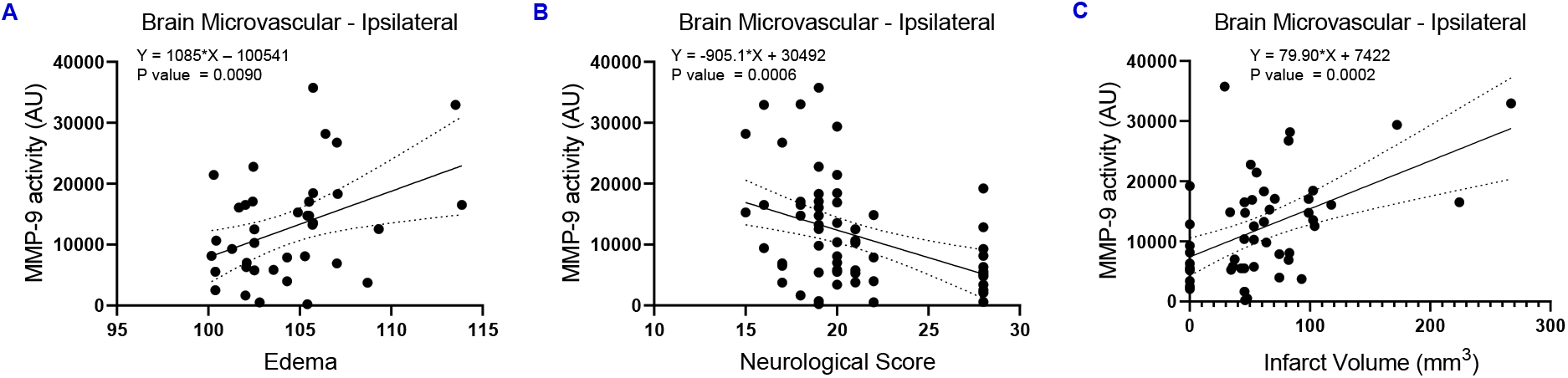
Correlation between cerebrovascular MMP-9 activity and edema, neurological function, and infarct volume. Graphs depicting MMP-9 activity (y-axis) observed in brain microvessels isolated from the ipsilateral hemisphere of rats post ischemic injury and at 24h reperfusion and the corresponding edema (A), neurological score (B), and infarct volume (mm^3^) (C) (x-axis). (A) *n* = 39 animals. (B) *n* = 62 animals. (C) *n* = 51 animals.

**Table 1.** Correlation of cerebrovascular MMP-9 activity, edema, neurological function, and infarct volume broken down by treatment condition.

| **Treatment** | **Variable 1** | **Variable 2** | **Slope (95% CI)** | **Y-intercept (95% CI)** | **R square** | **F (DFn, DFd)** | **P value** |
| --- | --- | --- | --- | --- | --- | --- | --- |
| **Sham** | MMP-9 | Edema | 1332 (-3545 to 6210) | -128554 (-620139 to 363032) | 0.1257 | 0.5753 (1,4) | 0.4904 |
| **Saline** | MMP-9 | Edema | 315.8 (-1913 to 2544) | -20151 (-252024 to 211722) | 0.02585 | 0.1327 (1,5) | 0.7305 |
| **rt-PA** | MMP-9 | Edema | 2057 (-13452 to 17565) | -200589 (-1847556 to 1446377) | 0.03278 | 0.1356 (1,4) | 0.7314 |
| **JNJ + rt-PA** | MMP-9 | Edema | 364.7 (-4117 to 4846) | -27919 (-496113 to 440274) | 0.005262 | 0.03703 (1,7) | 0.8529 |
| **BI + rt-PA** | MMP-9 | Edema | 3239 (-14986 to 21463) | -298202 (-2215616 to 1619212) | 0.04007 | 0.2087 (1,5) | 0.6669 |
| **JNJ + BI + rt-PA** | MMP-9 | Edema | 1758 (-1611 to 5127) | -169263 (-526384 to 187859) | 0.4790 | 2.759 (1,3) | 0.1953 |
| **Sham** | MMP-9 | Neurological Score | N/A | N/A | 1.000 | N/A | N/A |
| **Saline** | MMP-9 | Neurological Score | -1227 (-3875 to 1420) | 33166 (-19107 to 85440) | 0.08647 | 1.041 (1,11) | 0.3295 |
| **rt-PA** | MMP-9 | Neurological Score | -4352 (-8395 to -308.8) | 95085 (23485 to 166685) | 0.3971 | 5.929 (1,9) | 0.0377 * |
| **JNJ + rt-PA** | MMP-9 | Neurological Score | -199.7 (-5397 to 4998) | 15232 (-85960 to 116424) | 0.000732 2 | 0.007327 (1,10) | 0.9335 |
| **BI + rt-PA** | MMP-9 | Neurological Score | -8065 (-40547 to 24418) | 196810 (-451489 to 845109) | 0.04693 | 0.3447 (1,7) | 0.5756 |
| **JNJ + BI + rt-PA** | MMP-9 | Neurological Score | -2776 (-9838 to 4286) | 66959 (-61015 to 194933) | 0.3428 | 1.565 (1,3) | 0.2996 |
| **Sham** | MMP-9 | Infarct Volume (mm<sup>3</sup>) | N/A | N/A | 1.000 | N/A | N/A |
| **Saline** | MMP-9 | Infarct Volume (mm<sup>3</sup>) | 159.0 (6.578 to 311.5) | -1241 (-14652 to 12171) | 0.5898 | 7.190 (1,5) | 0.0437 * |
| **rt-PA** | MMP-9 | Infarct Volume (mm<sup>3</sup>) | 192.0 (-148.4 to 532.4) | 1001 (-25348 to 27351) | 0.2410 | 1.905 (1,6) | 0.2168 |
| **JNJ + rt-PA** | MMP-9 | Infarct Volume (mm<sup>3</sup>) | 74.29 (-120.7 to 269.3) | 6582 (-7886 to 21050) | 0.06721 | 0.7205 (1,10) | 0.4158 |
| **BI + rt-PA** | MMP-9 | Infarct Volume (mm<sup>3</sup>) | -63.12 (-1080 to 954.0) | 40964 (-50847 to 132775) | 0.003067 | 0.02154 (1,7) | 0.8875 |
| **JNJ + BI + rt-PA** | MMP-9 | Infarct Volume (mm<sup>3</sup>) | 92.04 (-41.75 to 225.8) | 7728 (-9874 to 25330) | 0.6151 | 4.793 (1,3) | 0.1163 |

\* Denotes a significant relationship between variable 1 and variable 2 following linear regression.

## Discussion

The present study evaluated whether pharmacological application of MMP-9 or/and LOX-1 inhibition in conjunction with rt-PA treatment in a clinically relevant experimental model of stroke improves the safety and efficiency of rt-PA treatment, thereby limiting brain damage development. Additionally, we evaluated the role of MMP-9 or/and LOX-1 inhibition on the cerebrovasculature and its barrier integrity in both an experimental stroke model as well as in vitro ischemia/reperfusion model utilizing primary HBMECs. To the best of our knowledge we have, for the first time, demonstrated that combination treatment with LOX-1 and MMP-9 inhibition conjugated with rt-PA not only improved the safety of delayed rt-PA treatment but significantly improved the stroke outcome in a thromboembolic model of stroke. In brief, delayed rt-PA treatment increased 1) murine hemorrhage and edema, 2) cerebrovascular MMP-9 activity and LOX-1 levels, 3) markers of HBMEC inflammation and activation, and 4) MMP-9 and/or LOX-1 inhibition differentially attenuated these responses (**Fig. 7**).

**Figure 7.**
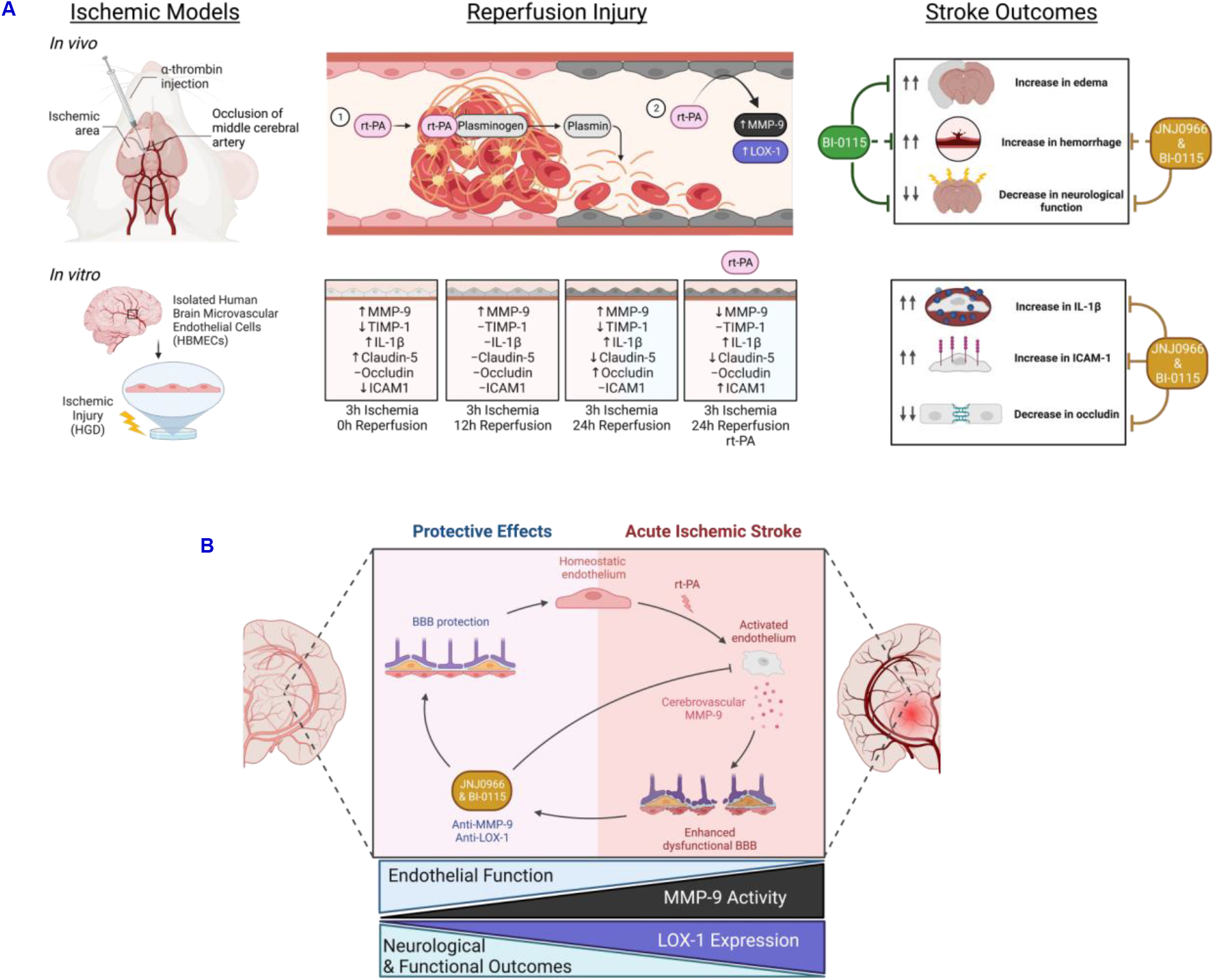
Schematic summary of cerebrovascular pathology during acute ischemic stroke in pre-clinical models. Illustrated is a simplified depiction of the contribution of recombinant tissue plasminogen activator (rt-PA) to vascular pathology during acute ischemia-reperfusion injury in both an in vivo and in vitro model of AIS. (A) rt-PA leads to increased LOX-1 expression and MMP-9 activity to endothelial dysfunction and the progression of ischemic stroke pathophysiology which can lead to endothelial vascular integrity disruption, oxidative stress, and inflammation. These contribute significantly to worsened AIS outcomes, and thus present an attractive therapeutic target. (B) These findings indicate that targeting MMP-9 and LOX-1 with novel selective compounds, JNJ0966 and BI-0115 respectively, improves stroke outcomes both in vivo and in vitro potentially by protecting against activated endothelium to promote a more homeostatic BBB following acute ischemia-reperfusion injury. rt-PA indicates recombinant tissue plasminogen activator; MMP-9, matrix metalloproteinase 9; LOX-1, lectin-like oxidized low-density lipoprotein receptor; HGD, hypoxia plus glucose deprivation; TIMP-1, tissue inhibitors of metalloproteinases 1; IL-1β, interleukin 1 beta; ICAM1, intercellular adhesion molecule 1; BBB, blood brain barrier.

In this study we sought to further investigate the pathogenic links between rt-PA, LOX-1, and MMP-9 in the context of AIS. Cerebral edema and hemorrhagic transformation are significant risks for administration of thrombolysis beyond the 4.5h therapeutic-window and have been linked to the vascular disruptors MMP-9 and LOX-1. Clinically, it has been reported that rt-PA therapy can increase the risk of hemorrhagic transformation by 6-8% ^42–44^ which can lead to an increased risk of worsened outcomes. In a separate meta-analysis, it has been reported that both endogenously produced tPA and rt-PA increases the risk of edema and disrupt the BBB in addition to increasing hemorrhage, culminating in increased mortality.^45^ Mechanistically, these increases are in part due to the off target effect of tPA which has been previously shown to activate MMP-9 ^46–48^ as well as increase parenchymal free radical production, increase leukocyte extravasation, and aggravate the microglia.^49^ The detrimental effects of rt-PA therapy depend highly on MMP-9 activation and breakdown of the BBB.^10^ In this study we demonstrated that the rt-PA treatment 4h post-stroke onset induced a significant increase in MMP-9 activity in isolated cerebral vessels (**Fig. 2H**). In addition, the delayed treatment with rt-PA resulted in worsened neurological function 24h after stroke onset (**Fig. 2G**). Similar results were previously observed in the same stroke model where delayed rt-PA therapy increased the risk of hemorrhagic transformation and increased the percentage of edema.^33^ This mimics clinical observations where delayed administration of rt-PA was provided to patients, thus indicating the relevance of this thrombolysis stroke model study.^10,50–52^

Due to the increase in MMP-9 activity at 6h post-thromboembolic stroke onset (**Figs. 1B & D**), we hypothesized that the most optimal effect on the safety of the treatment would include administration of the LOX-1 and MMP-9 inhibitors prior this timepoint. We, therefore, administered the inhibitors 3.5h after stroke onset, 30 min before the delayed rt-PA therapy. Administration of these selective inhibitors prior to rt-PA is advantageous as rt-PA treatment comes with an extensive list of contraindications ^53^ delaying treatment in patients when they first arrive in the clinic. Treatment with either or both LOX-1/MMP-9 inhibitors may not require this exclusionary step, making it possible to treat patients on arrival before rt-PA therapy, increasing translation to the clinic. Our observations of increased MMP-9 activity both within the parenchymal and isolated cerebrovascular tissues closely mimic the previously observed increase in homogenized brains of rats subjected to 1h of transient middle cerebral artery occlusion which demonstrated a trend increase in MMP-9 activity beginning at 4h post-injury.^54^ This is in opposition to a separate study which employed a model of permeant middle cerebral artery occlusion and observed the greatest brain and plasma MMP-9 activity at 24h post stroke.^55^ Together suggesting the importance of ischemic model selection, as the resolved occlusion and addition of reperfusion appears to increase MMP-9 activity both within the ipsilateral cerebrovascular and parenchymal tissue at 6h post-onset which was then decreased at 24h post-onset (**Figs. 1B-E**).

The complex role of the endothelium in maintaining a functional barrier is challenged by the enzymatic activity of MMP-9, which has been considered clinically as a marker of ischemic stroke and shown to actively exacerbate endothelial barrier permeability. It has been previously demonstrated in a murine model of intracerebral hemorrhage that MMP-9 was increased and co-localized with cerebral microvessels, ^56^ suggesting that the endothelium may play a significant role in the production of MMP-9 as observed in this study. Similar to our in vivo observations (**Fig. 2H**) we observed no increase in HBMEC derived MMP-9 activity following HGD3h/R24h injury (**Fig. 4B**) in the presence or absence of rt-PA (**Fig. 4C**). However, we observed a direct effect of both HGD3h/R24h and rt-PA on the alteration of HBMEC transcription (**Fig. 5**). Additionally, we observed that rt-PA lead to an increase in HBMEC-derived MMP-2 activity following HGD3h/R24h injury (**Fig. 4F**). We acknowledge that cultured cells do not identically match conditions made in vivo; however, to the best of our knowledge this is the first time HBMECs were exposed to ischemia reperfusion injury in the presence of rt-PA and further investigation into the temporal impact of the response of MMP-9 to these perturbations is warranted to elucidate potential mechanisms underlying this response. We also acknowledge that the complex pathophysiological cascade shown to occur during an ischemic stroke cannot be precisely modeled in an in vitro setting (i.e., lack of supplemental BBB components such as the CNS, mechanotransduction effect of blood flow, and peripheral immune cell responses). However, in vitro studies using human primary cells prompt the investigation of specific basic cellular and molecular mechanisms under conditions of hypoxia or oxygen and glucose deprivation plus reperfusion which is similar to what is observed during AIS.

At the level of the vasculature, LOX-1 has been shown to induce vascular smooth muscle and endothelial dysfunction leading to impaired nitric oxide relaxation and apoptosis.^57–63^ The overlapping activation of NF-κB both in the vascular smooth muscle and endothelium following LOX-1 stimulation has been shown to induce a positive feedback loop resulting in increased *ORL1* transcription and increased LOX-1 protein.^64^ This activation of NF-κB has also been linked to increased MMP-9 transcription ^65,66^ which has been highly implicated with worsening of ischemic injury pathogenesis.^38,67–69^ Previous efforts to address the role of LOX-1 and its subsequent detrimental role in vascular pathology support the potential beneficial effect of targeting LOX-1 to attenuate acute ischemic injury sequelae. Recently it has been shown that deletion of LOX-1 had protective effects in stroke-prone spontaneously hypertensive rats, which based on miRNA biomarker analysis, was proposed to be acting to attenuate LOX-1 induced dysfunction at the level of the BBB following cerebral ischemia.^70^ It has been previously found that mice overexpressing endothelial LOX-1 had increased stroke volumes and worsened neurological function at 24h following transient middle cerebral artery occlusion and treatment by LOX-1 silencing RNA had the reverse effect on these outcomes.^71^ Together suggesting that targeting of LOX-1 in the management and treatment of AIS may be a viable approach.

Based on the previously established connection between LOX-1 and MMP-9 we hypothesized that animals with an increased concentration of cerebrovascular LOX-1 protein would have increased MMP-9 activity following stroke which would lead to worsened outcomes (**Fig. 1F**). In this study we observed from 18 individual rats subjected to thromboembolic stroke that the ipsilateral cerebrovasculature LOX-1 protein levels were positively correlated with MMP-9 activity (**Fig. 1G**). To the best of our knowledge this is the first time that LOX-1 and MMP-9 levels and activity have been correlated within the cerebrovasculature following acute cerebral ischemic injury. While there has been no such previous connection established between these proteins within the cerebrovasculature under similar experimental conditions, there remains sufficient literature linking the LOX-1 to MMP-9 at the transcriptional and activation levels.

In this study, we overall, observed that the detrimental effects of rt-PA treatment were attenuated by LOX-1 and MMP-9 inhibition. LOX-1 inhibition in combination with rt-PA therapy reduced the infarct area as well as attenuated edema (**Figs. 2B-C**). These findings compliment a previous study which demonstrated that a deficiency or neutralization of LOX-1 reduced the infarct volume in a murine model of stroke at 24h post-injury onset following a 2h middle cerebral artery occlusion.^72^ Moreover, a separate study performed in rats on postnatal day 7 subjected to hypoxic-ischemic injury for 2h demonstrated that anti-LOX-1 antibody reduced edema as well as rescued tight junction proteins ZO-1 and occludin at 48 and 72h post-injury.^73^ Additionally, in this study we observed that targeting LOX-1 and MMP-9 separately and together showed a trend reduction in the prevalence and size of hemorrhagic events (**Fig. 2D**). These results confer with previously established reports demonstrating that inhibition of MMP-9 attenuates the increased risk of hemorrhagic transformation.^74^ However, to the best of our knowledge this is the first demonstration of the protective effects via LOX-1 inhibition conjugated with rt-PA on hemorrhagic transformation. This observed beneficial efficacy of LOX-1 and/or MMP-9 inhibition was also observed at the neurological level (**Figs. 2F-G**). It has been previously demonstrated in a rat model of cerebral ischemia/reperfusion injury that attenuating MMP-9 activity during the acute phase results in improved neurological function.^75^ However, unlike the 2013 study by Zhao et al, we did not observe an improvement in neurological function as determined by percent usage of injured forepaw in the rats treated with the MMP-9 inhibitor in conjunction with rt-PA (**Fig. 2F**). However, we observed a concomitant improvement in neurological function in rats treated with rt-PA and the LOX-1 inhibitor alone or in combination with the MMP-9 inhibitor. Moreover, we observed an increase in neurological score in rats treated with the selective LOX-1 inhibitor and rt-PA compared to rt-PA alone following thromboembolic stroke (**Fig. 2G**). Further suggesting that LOX-1 inhibition be improving neurological function. In this model, rt-PA has always had a negative impact on stroke outcome with delayed treatment, a phenomenon also seen in the clinic where the efficacy of rt-PA decreases with time. Here we show that in a model where the effectiveness of rt-PA is zero, we still see an improved outcome when combined with LOX-1 inhibition. This improvement in neurological function may be in part due to the anti-apoptotic effect previously observed by blocking LOX-1 in neonatal rats subjected to hypoxia ischemic injury.^73^

While the MMP-9 inhibitor had a more considerable influence on the MMP-9 activity than the LOX-1 inhibitor, both may be enough to increase the safety of rt-PA. Combined, they were still able to reduce MMP-9 activity. Intriguingly, we observed that not only did the selective MMP-9 inhibitor decrease MMP-9 activity, but it also decreased cerebrovascular LOX-1 protein levels at 24h post-injury (**Fig. 2I**). It has been previously demonstrated that LOX-1 activation can induce a positive feedback loop in which more LOX-1 protein is produced ^76^ and in line with this previous report we observed that the novel selective LOX-1 inhibitor reduced LOX-1 protein levels. However, contrary to what we expected, LOX-1 levels were not decreased in the cerebrovasculature of rats treated with the combination of both LOX-1 and MMP-9 inhibitors in addition to rt-PA which merits further investigation.

Based on our results in this study, we posit that the combination of LOX-1 and MMP-9 inhibitor is in part working at the level of the endothelium to reduce focal inflammation (IL-1β), endothelial activation (ICAM-1), and rescue tight junction expression (occludin) within the HBMEC to mitigate ischemia/reperfusion-induced decreases in cerebrovascular barrier integrity. Here we confirmed that MMP-9 is significantly associated with adverse functional outcomes such as increased edema, infarct volume, and decreased neurological score as well as that inhibition alone may be beneficial to prevent rt-PA induced risk of hemorrhagic transformation but will not improve stroke outcome as previously seen when minocycline was used in clinical trials. ^74^ BI-0115 is a selective LOX-1 inhibitor that prevents the binding of the oxLDL ligand and therefore blocks the signaling cascade and possibly the upregulation of MMP-9. In comparison, JNJ0966 is a direct MMP-9 inhibitor that prevents the activation of MMP-9 by binding allosterically to the protease. The study presents the road map to future vascular based strategies. The focus on extending the window of opportunity for rt-PA treatment will have immediate effects for millions of patients and healthcare systems worldwide.

## Supplemental Tale and Figure Legends

**Supplemental Table 1. Correlation of extracellular HBMEC MMP-9 and MMP-2 activity with HBMEC mRNA levels following HGD/R treated with either vehicle, rt-PA, or JNJ + BI + rt-PA.**

* Denotes a significant correlation between variable 1 and variable 2 following Pearson’s correlation.

**Supplemental Table 2. Sense and anti-sense strands of primers used for qRT-PCR.**

**Supplemental Figure 1. Spatiotemporal brain microvascular and parenchymal LOX-1 expression following thromboembolic stroke.** (A & D) Representative western blots illustrating the band migration for LOX-1 (32kDa) and the loading control B-Actin (42kDa), in isolated brain microvascular and parenchymal tissues from the ipsilateral and contralateral hemispheres following either sham or thromboembolic stroke at 3, 6, and 24h. Bar graphs depict LOX-1 protein levels within isolated microvessels (B-C) and parenchymal tissue (E-F) from the ipsilateral (B & E) and contralateral (C & F) hemispheres relative to injury by densiometric analysis and expressed as arbitrary units (AU) normalized to B-Actin *±* SD. *n* = 3-7 animals (data are mean ± SD).

**Supplemental Figure 2. Impact of rt-PA +/− LOX-1 and/or MMP-9 inhibition following thromboembolic stroke on contralateral brain microvascular MMP-9 activity and LOX-1 expression.** (A) Bar graphs depict MMP-9 activity (A) and LOX-1 protein levels (B) within isolated microvessels from the ipsilateral hemispheres relative to injury by inverse densiometric (A) and densiometric (B) analysis and expressed as arbitrary units (AU) (A) and arbitrary units (AU) normalized to B-Actin (B) *±* SD. *n* = 5-8 animals (data are mean ± SD).

**Supplemental Figure 3. Impact of rt-PA +/− LOX-1 and/or MMP-9 inhibition following HGD/R on HBMEC H2O2 production, SOD1, and ZO-1 expression.** (A-B) Bar graphs depict extracellular hydrogen peroxide levels following HGD/R (3h/24) exposure relative to normoxia (A) and following treatment with rt-PA *±* MMP-9 + LOX-1 inhibitors as *μ*mol/L. (C-D) Bar graphs depict extracellular hydrogen peroxide levels following HGD/R (3h/24) exposure relative to normoxia (A) and following treatment with rt-PA *±* MMP-9 + LOX-1 inhibitors as *μ*mol/L. Bar graphs depict qRT-PCR mediated quantitation of SOD1 mRNA present within HBMECs exposed to either normoxia or HGD/R at 3h/24h (C) or HGD/R (3h/24h) and treated with vehicle, rt-PA, or rt-PA + MMP-9 + LOX-1 inhibition (D) normalized to the housekeeping gene (GAPDH) and expressed as 2^−ΔΔ*ct*^. Samples were run in duplicate. (E) Bar graph depicts qRT-PCR mediated quantitation of ZO-1 mRNA present within HBMECs exposed to either normoxia or HGD at 3h followed by HGD/R for 3h/12h and 3h/24h normalized to the housekeeping gene (GAPDH) and expressed as 2^−ΔΔ*ct*^. Samples were run in duplicate. *n* = 4-12 (data are mean ± SD).

## Acknowledgments

We are grateful to Lund University Bioimaging Centre for their assistance and providence of the MRI facility. We also thank Dr Emmanuel Barbier at Grenoble Institute of Neurosciences for sharing the method for pCASL.

## Sources of Founding

This work was supported by the Brain Foundation, Crafoord Foundation, Franke and Margareta Bergqvist Foundation, Neuro Foundation, Olle Engqvist Foundation, Swedish Heart-Lung Foundation, Swedish Stroke Foundation, and Thure Carlsson Foundation. The American Heart Association and The UA COMP Valley Research Partnership Award.

## Notes

### Competing Interest Statement

The authors have declared no competing interest.

